# Source amplitude increases with body mass across avian genera

**DOI:** 10.1101/2024.10.31.621325

**Authors:** Morgan A. Ziegenhorn, Richard B. Lanctot, Stephen C. Brown, Sarah T. Saalfeld, Paul A. Smith, Nicolas Lecomte

## Abstract

Amplitude, or intensity, of sound is a fundamental characteristic of inter acoustic communication, with relevance in many scientific fields. The amplitude of an animal’s acoustic signal at its source (“source amplitude”) may be particularly relevant in the field of acoustic allometry, where relationships between species’ physical and acoustic features (e.g., dominant frequency) have been well-established across taxa. However, despite their potential scientific value, records and studies of source amplitudes remain remarkably scarce for avian species. Here we present novel estimates of source amplitude (range, mean, and median) for 17 species of Arctic-breeding birds, derived from measurements made in Utqiaġvik, Alaska during June 2024. We found a strong positive correlation between body mass and source amplitude in this data via a Markov Chain Monte Carlo multivariate generalized linear mixed model (MCMCglmm). Both phylogeny and individual identity were important random effects in this model. In contrast, random effects from environmental factors and measurement characteristics were minimal. Our work represents the first model describing the relationship between source amplitude and allometry across avian genera. We hope that this study will spur other investigations into avian source amplitude and its relationship to morphological and life history features for species in the Arctic and elsewhere.

## Introduction

Communication via sound is ubiquitous in avian species. Many types of vocalizations, including contact calls (Meaux et al. 2023; Sharp et al. 2005), songs and display calls (DiSciullo et al. 2024; Favaro, Ozella, and Pessani 2014; Sebastianelli, Blumstein, and Kirschel 2022; Sierro, de Kort, and Hartley 2023), alarm calls (Leger and Nelson 1982; Magrath, Pitcher, and Gardner 2007), and territory defense calls (Morton and Stutchbury 2012; Nowicki, Searcy, and Hughes 1998), among others, serve important functions in bird communities. Over the past century, bioacoustic research has focused on the description, documentation, and analysis of these vocalizations, offering critical insights for species’ conservation and behavioral ecology. One aspect of this research, acoustic allometry, explores the relationship between body size and features of animal vocalizations, as well as the plasticity of these complex signaling traits in different ecological contexts. Prior studies have found allometric relationships between frequency-related components of sound (e.g., dominant frequency, signal bandwidth, fundamental frequency) across a variety of taxa (e.g., (Bowling et al. 2017; Fletcher 2004; Garcia et al. 2017; McCracken and Sheldon 1997; Pélabon et al. 2014)). Deviations from these allometric relationships have been linked to factors such as sexual selection (Garcia and Ravignani 2020; Tonini et al. 2020), presence of acoustic competitors (Díaz-Torres et al. 2022; Memet, Farrell, and Mahadevan 2022), anthropogenic noise sources (Augusto-Alves, Dena, and Toledo 2021), and terrain (Riondato et al. 2021).

While relationships between allometric features and vocal frequency measures are common, source level (for this paper, “source amplitude”)—a key aspect of acoustic communication—has been largely overlooked. Amplitude is necessarily crucial to acoustic communication as it directly constrains the distance at which an individual can be perceived by another. In addition to this, source amplitude is thought to potentially be an honest signal of body size (Gray 1997) and aggressive intent (e.g., (Akçay, Beck, and Sewall 2020; Searcy, Anderson, and Nowicki 2006)). Source amplitude often also serves as a crucial benchmark for understanding behavioral shifts such as the Lombard effect, where amplitude is increased in response to increased background noise, often from anthropogenic sources. Such effects have been demonstrated for taxa such as marine mammals (Erbe et al. 2016; Guazzo et al. 2020; Holt, Noren, and Emmons 2011), bats (Lewanzik and Goerlitz 2018; Surlykke and Kalko 2008), primates (Hotchkin, Parks, and Weiss 2013; Schopf, Schmidt, and Zimmermann 2016), and a few birds (H. Brumm and Ritschard 2011; Henrik Brumm 2004; Dorado-Correa, Zollinger, and Brumm 2018; Reichard and Anderson 2015). Increasing source amplitude to cope with shifting background noise may incur metabolic or social costs (e.g., increased attacks by territorial conspecifics), while an inability to increase amplitude will necessarily result in decreased effective communication space (or active space of the signal) (Guazzo et al. 2020). For these reasons, documenting species’ amplitude ranges and describing relationships to body mass and other factors form a key basis for monitoring shifts in the face of increased anthropogenic or environmental pressures.

The scarcity of data on source amplitude among the 10,000+ bird species is perhaps due to the challenges of measuring it accurately in the field, where distance from the subject, environmental obstacles such as trees, and a lack of calibrated recording equipment can heavily influence results. Only one study to date has documented source amplitude scaling with body mass across species (17 songbirds, and the domestic chicken)(Brackenbury 1979). However, this study did not include consideration of a potential phylogenetic signal, which may also play a role in the degree of scaling between mass and acoustic properties of communication. Rodriguez et al. (2015) suggested that the degree of allometric scaling between body size and frequency parameters in birds, tree frogs, tree crickets, and crickets depended upon taxa-related differences in mating systems (i.e., whether mate preferences favored ‘stabilizing’ signal trait values, or ‘extreme’ values). Notably, this study mentioned the lack of available data on source amplitude as a weakness and area of future work. In birds, phylogeny has also been shown to play a significant role in the strength of allometric relationships between frequency parameters and body size for some species (Friis et al. 2021; Laiolo, Caprio, and Rolando 2003; I. Torres, Lopez, and de Araújo 2017; I. M. D. Torres, Barreiros, and de Araújo 2020). The role of phylogeny in relation to source amplitude is nearly unexplored, though Brumm (2009) suggested that louder signals in two passerine species might elicit stronger aggression responses, hinting at a relationship to species’ life history strategies, which may be phylogenetically conserved. A recent study of several other passerines species has also found significant phylogenetic signals in both the morphological and behavioral components of song structure (Mejías et al. 2020). Investigating avian source amplitude in an allometric context may provide similarly significant insights into life history strategies and species’ relationships to their ecological niches.

In this study, we addressed the significant research gap in source amplitude documentation across 17 avian species. We also quantified the allometric relationship between source amplitude and body size for these species, while considering the relative impact of phylogeny, individual identity, and environmental factors on this relationship. Given previous evidence of phylogenetic and allometric components to features of species’ vocalizations, we predicted a positive allometric relationship between body size and source amplitude, with a strong phylogenetic signal. To test our prediction, we used source amplitude estimates from 17 avian species recorded vocalizing during the early nesting period near Utqiaġvik, Alaska.

## Materials and Methods

### Data collection

Amplitude measurements of avian vocalizations were made in a variety of tundra habitats in and around Utqiaġvik, Alaska between June 2-22, 2024. The relatively flat tundra habitats of Alaska’s North Slope host numerous migratory bird species, including waterfowl, seabirds, shorebirds, and landbirds (Johnson and Herter 1989). The tundra is uniquely advantageous for amplitude measurement, as background noise (aside from wind) is minimal and obstacles (e.g., trees, large rocks) that might scatter sound are virtually nonexistent. In addition, the relative ease of approaching arctic-breeding birds and the lack of obstacles allows observers to visually detect breeding birds at a relatively close range with minimal disturbance. The birds recorded were primarily shorebirds, though vocalizations from several waterfowl, seabirds, and passerines were also measured (see Table 1 for the full species list).

**Table 1.**
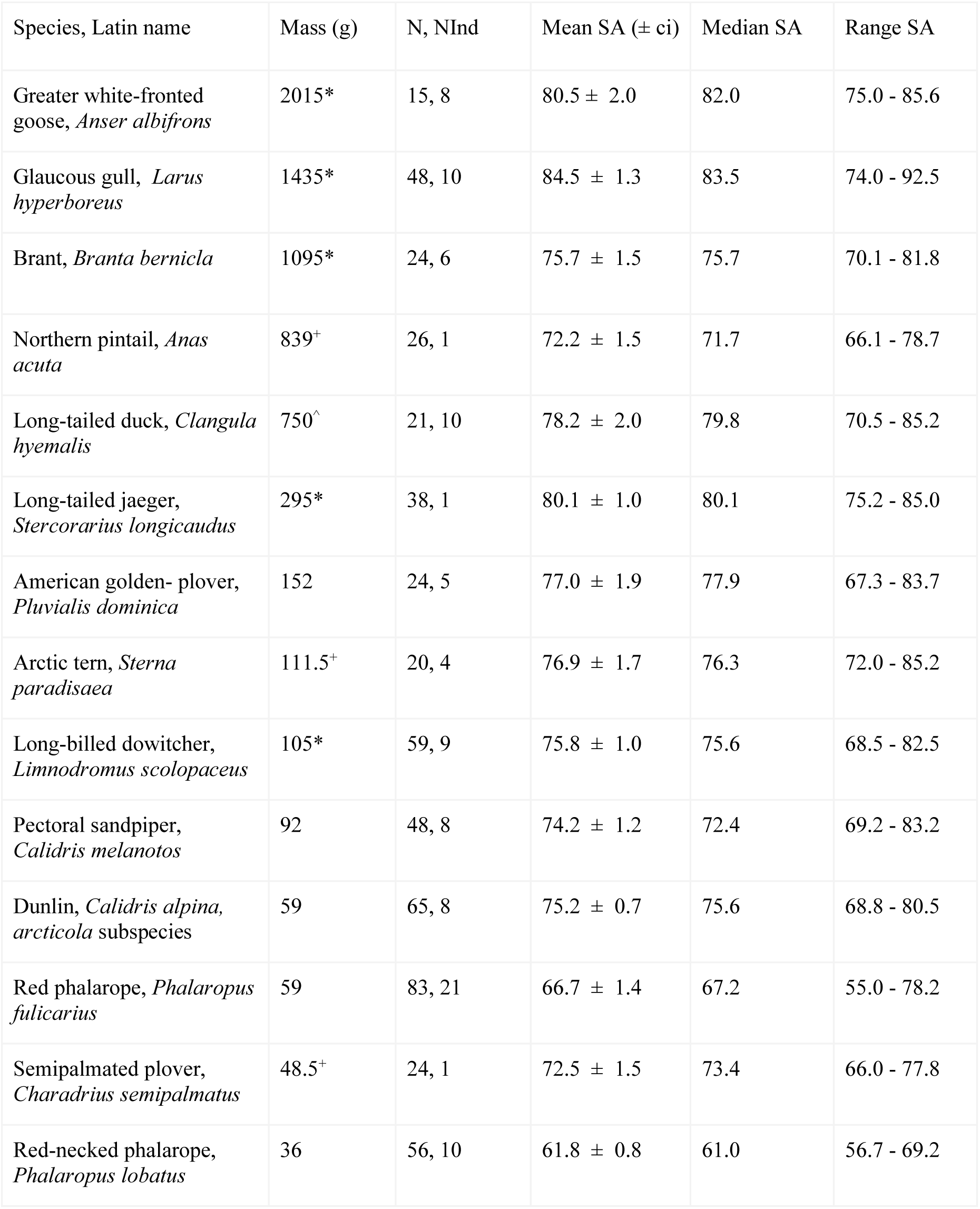
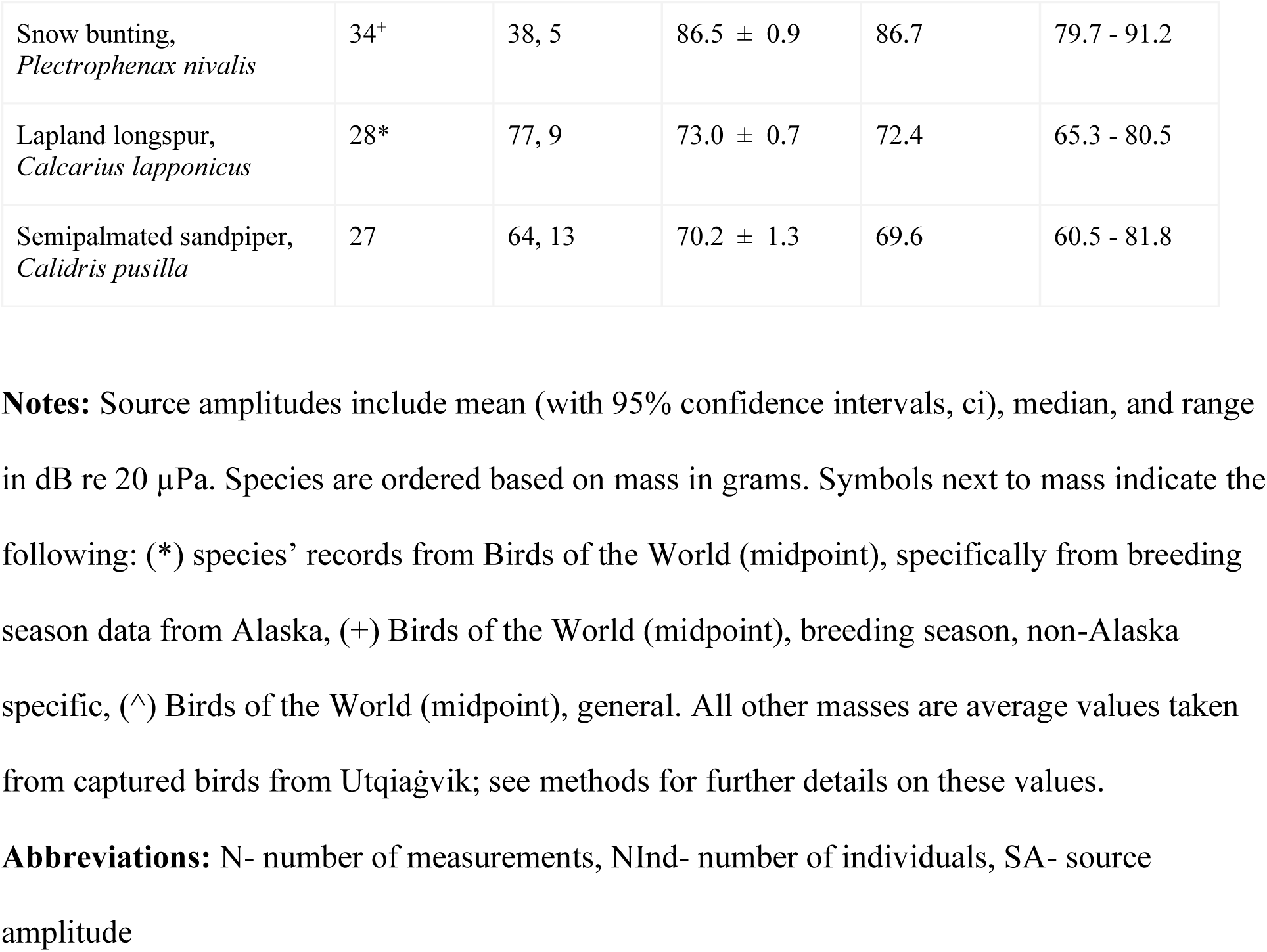
Source amplitude estimates from field measurements of 17 Arctic-breeding birds from Utqiaġvik, Alaska, June 2024.

We used a LATNEX sound pressure level (SPL) meter (SM-130DB), which instantaneously measured amplitude of sound in the environment as SPL in dB (fast time integration of 125 ms, A-weighted, dB re 20 µPa) and follows the most recent sound level meter standards (Commission 2003). A windscreen ball (65 mm diameter) was used on the SPL meter at all times to mitigate background noise. In addition, measurements were not made on days when wind speed exceeded 32 kph and were generally avoided on days when wind speed exceeded 24 kph. When wind was noticeable during measurement, care was taken to position the measurer’s body between the SPL meter and the wind to provide additional cover, though never between the SPL meter and the vocalizing bird. Measurements were not taken during periods when environmental noise might have affected readouts, i.e., when noise from nearby roads, plane traffic, or other birds was noticeable. At all other times, background noise on the tundra was quite minimal.

Measurements were taken by positioning the SPL meter directly towards a vocalizing bird that was also facing towards the SPL meter. In all cases, no obstacles were between the vocalizing bird and the SPL meter during measurements, minimizing sound absorption and scattering of sound waves by objects in the environment. The maximum amplitude for each vocalization produced was recorded; multiple measurements were often taken from one individual as it produced a variety of vocalizations. Measurements were taken regardless of sex and vocalization type to better describe the full range of amplitudes used by a given species. For the same reason, measurements were made in a wide variety of behavioral contexts, including mating displays, nest defense, and territory defense.

Distance from the bird was estimated visually at close range. Given the short distances involved (generally < 20 meters), and the small sizes of the birds, it was not practical to estimate distances with a rangefinder. Distance was not measured with paces or a tape measure as this would have disturbed the bird. We examined the potential impact of distance from the bird by including distance as a random effect in the model exploring the relationship between body size and source amplitude.

As mass was not recordable in the field, we used the average mass for shorebirds captured at Utqiaġvik, Alaska from 2003-2024 during the same period (June 1-22) and general location where the recordings were obtained. Adults were captured during pre-laying or at nest sites and weighed using a pesola scale accurate to the nearest 2 grams. We used the average mass for female phalaropes and male pectoral sandpipers because they are the vocal sex and hence were the subject of most (if not all) of our recordings. These values were used for all shorebird species except long-billed dowitcher and semipalmated plover, due to insufficient sample size (n = 3 and n = 0, respectively). For all other species, mass was calculated by species as the midpoint of the range provided by Birds of the World (Billerman et al. 2022). Specific mass ranges from breeding birds in Alaska were used whenever possible. Otherwise, ranges from breeding birds elsewhere were used. For long-tailed ducks, no breeding specific range was available, so the overall range was used. Where ranges differed for males and females, we first took the midpoint for each sex and then averaged those numbers (Table 1). Mass was log-transformed (base 10) for all further analyses to make the distribution of values more normal. Our environmental variables included wind speed and direction, temperature, and humidity. Hourly values of these variables at the Barrow airport (approximately 5-10 km away from measurement locations) were downloaded from the NOAA National Center for Environmental Information’s Climate Data Online service (https://www.ncei.noaa.gov/cdo-web/datasets/GHCND/stations/GHCND:USW00027502/detail). The hour-specific values of these variables were associated with each recording based on the time of the recording.

### Data analysis

Source amplitude was calculated from all collected amplitude measurements using the spherical spreading model for sound propagation from a point source:

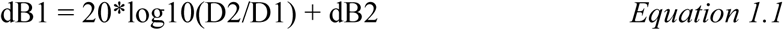

where dB1 is amplitude at distance D1 from the source, and dB2 is amplitude at distance D2. With a standard reference distance of 1 meter, D1 = 1 and this simplifies to:

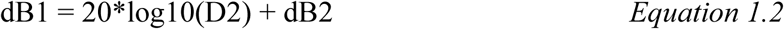

where D2 is our measurement distance from the bird.

For each species, mean, median, and range of source amplitudes were calculated.

Multiple measurements from the same individual were retained for analyses to better describe the overall variation in amplitude. As the number of measurements was uneven across species, we conducted a simple linear regression across all species that investigated the relationship between the number of measurements for a given species and that species range of source amplitude values.

We evaluated the relationship between body mass and source amplitude using a Markov Chain Monte Carlo multivariate generalized linear mixed model (MCMCglmm in R, (Hadfield 2010)). This model was constructed with source amplitude value as the response variable, and species’ body mass (log scale) as a fixed effect, and all other considered variables that might influence this relationship as random effects. Recording-specific variables included species identity, individual identity (as multiple measurements were taken from individuals), measurement distance, and measurement date (day of month), while environmental variables included wind speed and direction, temperature, and humidity. Environmental conditions (wind, temperature, and humidity) were included due to their potential effects on how sound propagates away from its source. Potential effects from the genetic relatedness among species was examined using a phylogenetic distance matrix as an additional input. The phylogenetic distance matrix was computed using classification hierarchies from the NCBI taxonomy database, extracted and compiled using the taxize package in R (Chamberlain and Szöcs 2013). This matrix included all species with source amplitude measurements. Prior to modeling, pairwise correlations between the above variables were computed, with an acceptable threshold of 0.6; no correlations were above this threshold so all variables were retained in the model construction.

Our MCMCglmm used weak priors for all random effects, with 600,000 iterations, burnin of 5,000, and thinning of 50. These values were increased from the default for iteration, burnin, and thinning to ensure model convergence. We reported the slope, credible intervals, and p-value for the regression line between body mass and source amplitude. The influence of all random effects was described using lambda, i.e., the ratio of the posterior values for a given random component to the posterior values of residual units (Garamszegi 2014). Lambda ranges from 0 to 1, where a high (>0.5) lambda indicates a strong effect, while a lambda <0.5 indicates a weak relationship to the response variable (source amplitude, in this case). All analyses and visualizations were performed using the R statistical platform (version 4.2.3, (R Core 2021)).

## Results

Our final analyses included 730 estimates of source amplitude from 129 individuals of 17 species (Table 1, Fig. 1). Across species, the loudest estimated source amplitude was 92.5 dB (glaucous gull), while the quietest estimate was 55.0 dB (red phalarope) (Fig. 1, Table 1).

**Figure 1.**
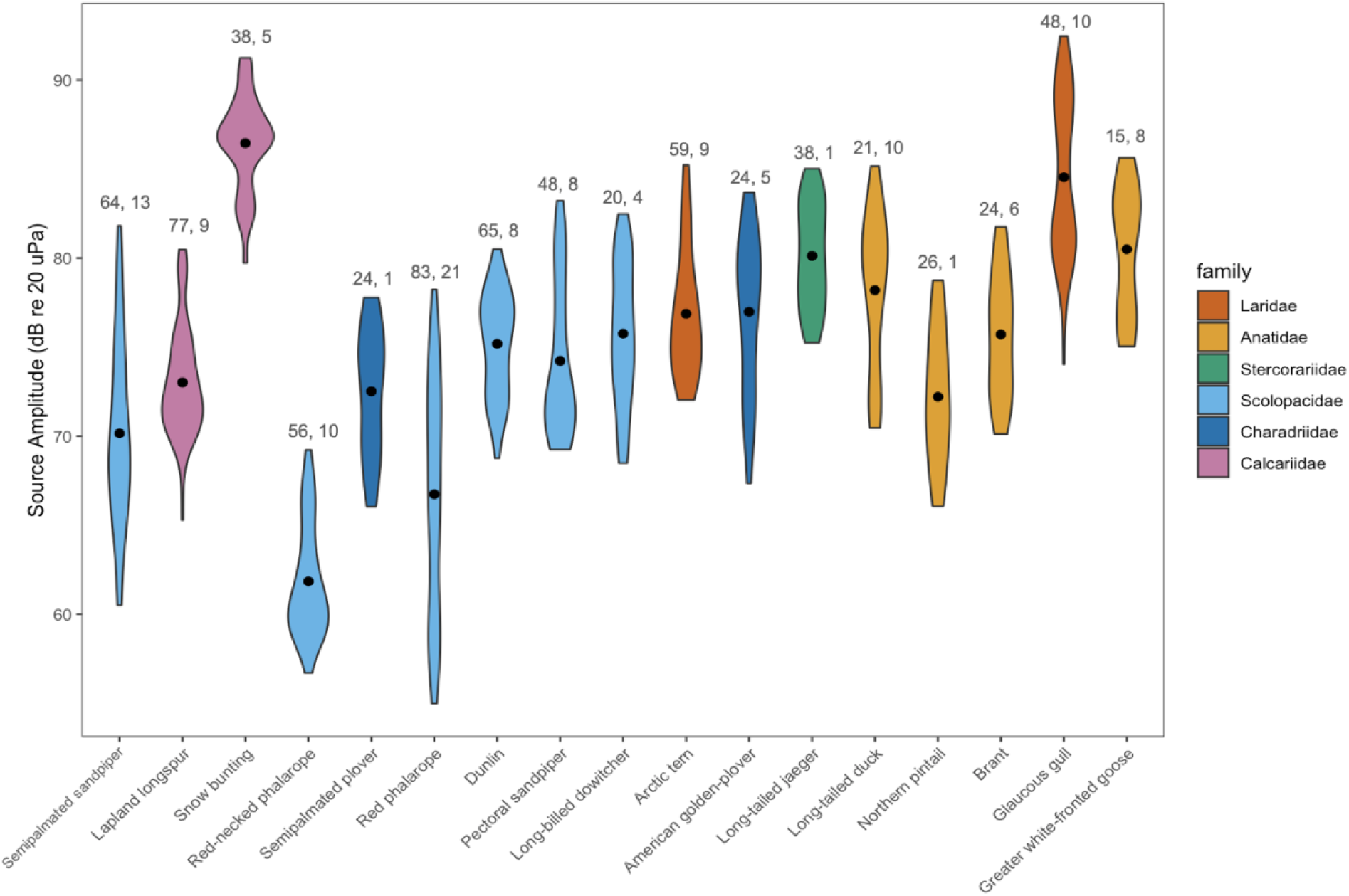
Source amplitudes for 17 species of arctic-breeding birds as measured in Utqiaġvik, Alaska in June 2024. Violin plots show variation in source amplitude estimates by species, with mean source amplitude shown as a black dot. Species are ordered based on body mass in grams (increases from left to right). Number of measurements and number of individuals are listed above each species. Distributions are colored by phylogenetic family.

Amplitude range was greatest for red phalaropes (55.0 - 78.2 dB) and semipalmated sandpipers (60.5 - 81.8 dB), and smallest for long-tailed jaegers (75.2 - 85.0 dB). Mean and median amplitudes were highest for snow buntings (85.1 and 86.5 dB, respectively) and lowest for red-necked phalaropes (61.8 and 61.0 dB, respectively). Assuming our distance estimates were accurate within +/- 0.5 meters at 3 meters, and 5 meters at 50 meters (the range of distances included in our analysis), our maximum inaccuracy in amplitude would be 1.1 dB at 3 meters and 0.9 dB at 50 meters, via Eq. 1.2. The mean distance (±SD) between the bird and the recorder was 22.6 ± 12.1 m. At this distance, an error in estimation of 1 meter would amount to an inaccuracy of 0.4 dB, while a maximum error of 5 meters would increase inaccuracy to 2.2 dB. We found no significant relationship between the range in source amplitude and number of measurements recorded for a given species (p-value = 0.655, adjusted R2 = −0.0202).

We found a statistically significant relationship between source amplitude and species’ body mass in our model (slope: 6.847, credible interval, ci: 0.420 - 13.6, p-value: 0.0418; Fig. 2). Importance of random effects on this relationship was variable, with both species (i.e., phylogeny) and the number of individual recordings having strong effects (i.e., lambda = 0.931 and 0.811, respectively). All other random effects (measurement distance, measurement date, wind speed, wind direction, temperature, and humidity) were less important, with lambda values below 0.5. Measurement distance came closest to this cutoff, with a lambda value of 0.42, while the relative importance of temperature, wind speed, and wind direction were particularly low (lambda less than 0.1 in all cases, Table 2).

**Figure 2.**
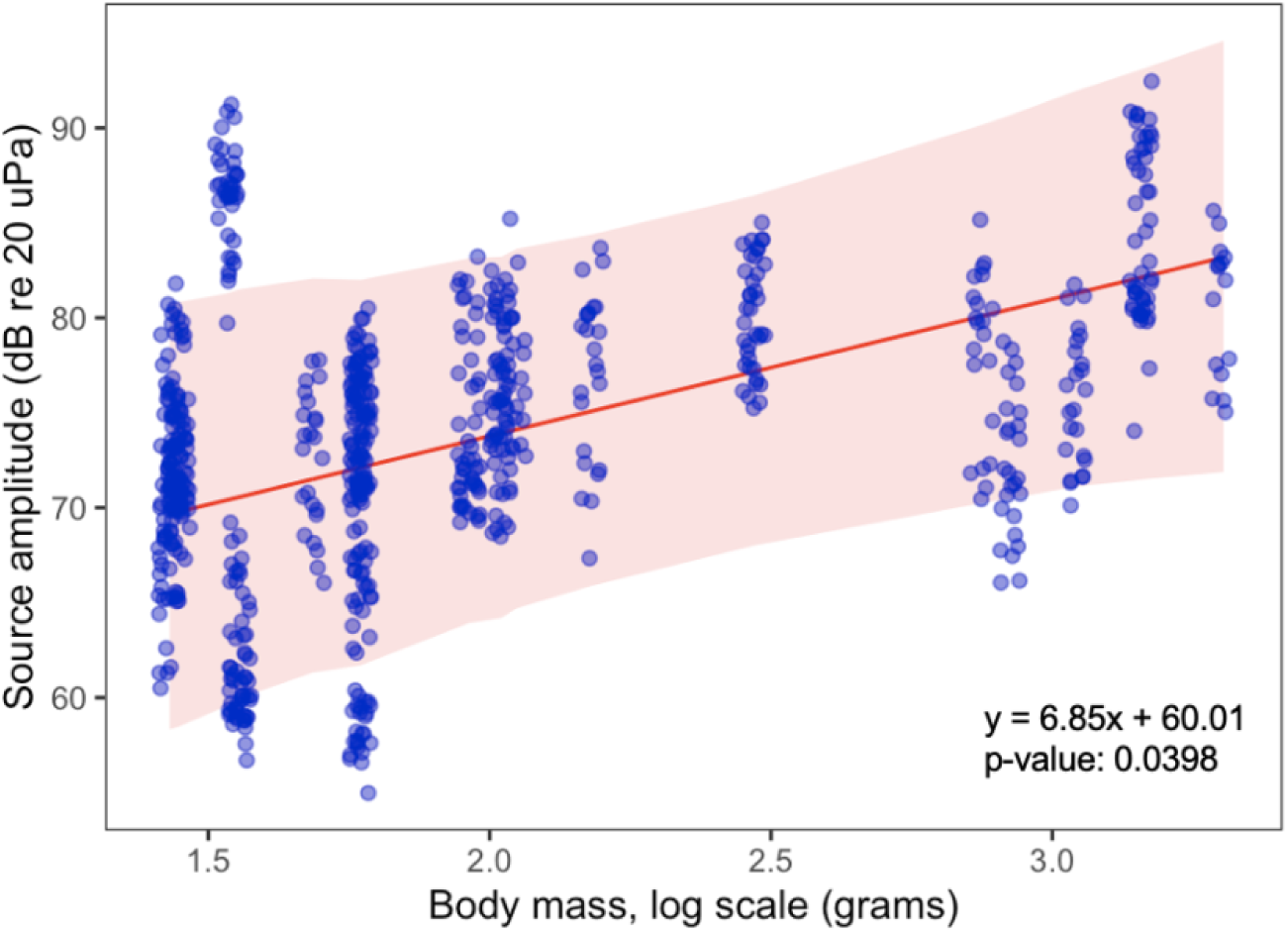
Model results. Modeled relationship (via MCMCglmm) between body mass and source amplitudes among 17 species of arctic-breeding birds as measured in Utqiaġvik, Alaska in June 2024. Blue circles indicate the data used to build the model. Predicted values are shown as a red line, with shading representing 95% credible intervals. Equation for the regression line and p-value are shown in black in the bottom right of the plot.

**Table 2.**
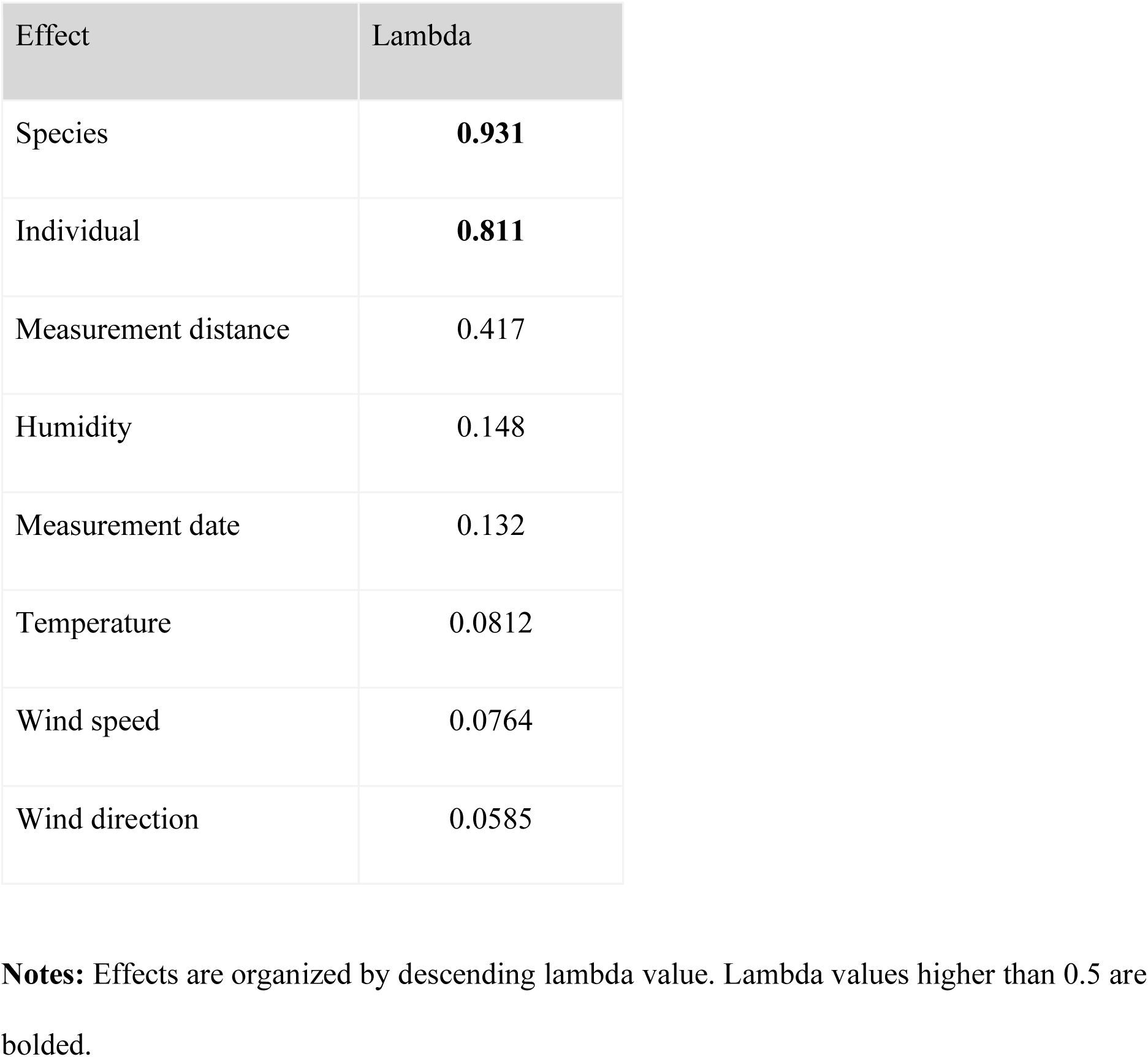
Lambda values for each random effect included in the MCMCglmm model explaining the relationship between body mass and source amplitude across 17 species of Arctic-breeding birds.

## Discussion

In this study, we produced the first record of range, average, and median values of estimated source amplitude for 17 species of Arctic-breeding birds, including the first source amplitudes for any species of shorebird or seabird. Our targeted taxa included waterfowl, seabirds, shorebirds, and passerines, with shorebirds as the most represented group (eight species). Our reported ranges for source amplitude were variable amongst species but are generally in line with previously published source levels (anywhere from 55 to 95 dB, generally, though notably no records are from close relatives)(Anderson et al. 2008; Brackenbury 1979;

Henrik Brumm 2004; Dorado-Correa, Zollinger, and Brumm 2018; Zollinger, Goller, and Brumm 2011). We also showed that collecting more measurements from the species in our study did not result in a wider range of measured source amplitudes for a given species. This suggests that uneven sampling amongst species did not affect our results.

There may be several reasons for species’ to produce sounds within a wide range of amplitudes. Behavioral state, vocalization type, and social context (H. Brumm and Ritschard 2011) are the most likely causes of this variability. Although we did not differentiate songs from calls during field recording, this information would likely provide further insight into source amplitude variation for our species. This may be particularly true for sexually dimorphic, polygynous species (e.g., pectoral sandpiper, red phalarope) for which mass and vocal behavior differ significantly between males and females. Previous studies have also noted that environmental variability can shift source amplitudes, perhaps most notably via the Lombard effect (Henrik Brumm and Slabbekoorn 2005; Lowry, Lill, and Wong 2012). As we controlled for environmental variability as much as possible, and made measurements in the relatively quiet tundra, it is possible and perhaps even likely that we have not captured the full range of source amplitudes that our study species can produce. Investigating the included species in noisier contexts, as well as including the effects of behavior, vocalization type, and sex in greater detail, are all valuable avenues for future study.

Apart from its relevance for animal behavior, source amplitude has a variety of applications in passive acoustic monitoring (PAM) in birds and other vocal taxa, which is increasing in scope and popularity. For example, source amplitude is used by PAM practitioners to localize animals (Putland et al. 2018; Tolkova and Klinck 2022) and understand the effective survey range of PAM recorders (e.g., (Winiarska, Szymański, and Osiejuk 2024)). The latter is important when using PAM for density estimation (Barlow et al. 2021; Pérez-Granados and Traba 2021), a common goal of many PAM conservation efforts. Based on our data and Eq. 1.2, an average red-necked phalarope with a source amplitude of 61.8 dB would have a general maximum detection range of 69 meters, assuming that any sound quieter than 25 dB would be undetectable by the recorder at 1 meter in ideal conditions. In contrast, an average snow bunting with a source amplitude of 85.1 dB would have a general maximum detection range of slightly more than 1 kilometer. While these are extremes, they demonstrate the extent to which inter and intra-specific variability in source amplitude has significant implications for PAM research.

Perhaps most crucially in this study, we provide a novel investigation into the potential allometric relationship between source amplitude and body mass amongst avian species while acknowledging the potential random contributions of several other factors. Our model demonstrated that most random effects (measurement distance, temperature, humidity, wind direction, wind speed) had very minimal effects on source amplitudes (Table 2). This is important evidence that recording opportunistically under somewhat controlled environmental conditions resulted in reliable data, despite having methodology that was relatively lax regarding these potential confounding factors. This is particularly important with regards to distance. Even if distance plays a limited role in source amplitude error, the maximum potential error (e.g., 2.2 dB at 22.6 meters with an inaccuracy of 5 meters) is relatively minimal in comparison to the range of source amplitudes produced by a given species. The analysis also indicated that measurement date, which may relate to behavior in terms of breeding phenology, did not introduce important variability into our data.

Our results suggest that an allometric relationship, with a strong phylogenetic signal, does exist between body size and source amplitude for these diverse arctic-breeding avian species.

The simplest explanation for the relationship to body size is that size directly limits the physical mechanisms of sound production in the syrinx (Brackenbury 1979), and hence limits the maximum possible source amplitudes. In our study, passerines prove the most notable exception to this, as source amplitudes were higher than would be expected for their size (Fig. 1-2). This may be related to the presence of intrinsic muscles in the syrinx that provide additional strength to the sound production system (Brackenbury 1979) that may not be present in the other clades represented (e.g., Charadiiformes, (Brown and Ward 1990)). These differences in sound production mechanisms among families may partially explain the strength of the phylogenetic signal. Amplitude ranges and maximums may also be behaviorally constrained by the way sexual selection within a species has influenced call/song production, as has been noted as an explanation for the body mass-frequency relationship in several cases (e.g., (Augusto-Alves, Dena, and Toledo 2021; Garcia and Ravignani 2020; Memet, Farrell, and Mahadevan 2022)). If more closely related species have more similar mating systems, then this could also explain the importance of phylogeny in our model. As more records of source amplitude become available, it will be possible to investigate this relationship in greater detail and derive important insights into how and why deviations from the allometric relationship between body mass and source amplitude occur (e.g., (Tonini et al. 2020)).

Individual variation in source amplitude was the second strongest random effect after species in our MCMCglmm (lambda 0.931 and 0.811, respectively). The strength of this relationship was robust to removal of species with less than 5 individuals from the model (lambda 0.819, in this case). The importance of this factor has been noted in several other studies (e.g., (H. Brumm and Ritschard 2011; Patricelli, Dantzker, and Bradbury 2008; Podos and Cohn-Haft 2019)). How individual variation in source amplitude values is related to individual body size is worthy of further study, especially in the context of amplitude as an honest signal of mate fitness. Studies of this type are currently quite rare; though Brumm (2009) found no intraspecific relationship between body mass and source amplitude in zebra finches, *Taeniopygia guttata*, or common nightingales, *Luscinia megarhynchos*. Instead, environmental noise and presence of other singers played a more important role in determining maximum singing volume. Similarly, the individual variation seen in our data may also highlight the plasticity of source amplitude, suggesting that context plays a dominant role in determining source amplitude moment-to-moment within a physically constrained range of potential amplitudes. Further study of this plasticity would ideally consider vocalization type, as this is unaccounted for in our case.

Documenting and investigating species’ source amplitudes provides key insights that can be applied in ecological and behavioral research, as well as conservation and monitoring programs. In turn, incorporating source amplitude into the field of acoustic allometry provides a novel avenue for further research into the complexity of its relationship to body size, as well as where and why deviations from that relationship may exist. As species’ environments get noisier and conserving them becomes both more crucial and more complicated, it remains important to document and understand such relationships. We hope this research will provide a basis for understanding changes in these behaviors, especially with increasing anthropogenic changes occurring across the Arctic.

